# Development of Collagenous Filaments with Tuneable Mechanical Properties Using a 3D Bioprinter and Molecular Crowding

**DOI:** 10.1101/2025.10.29.684883

**Authors:** Alejandro Rossello, Zixie Liang, Zihan Zhou, Xiangyu Gong, Ryan Nguyen, Seyma Nayir Jordan, Michael Mak

## Abstract

Engineering collagen constructs that replicate the mechanical strength and biological functionality of load-bearing tissues like tendon remains a significant challenge. This study establishes a scalable and accessible biofabrication method that combines the precision of a commercial 3D bioprinter with the physicochemical principles of molecular crowding. Using an acidic collagen solution in combination with a hypertonic polyethylene glycol (PEG) bath, we systematically investigate how modulating pulling speed and nozzle diameter dictates the final filament properties. Our results demonstrate that faster pulling speeds and smaller nozzle diameters produce thinner, more densely packed filaments. High-resolution scanning electron microscopy (SEM) and polarised light microscopy reveal a tuneable transition in surface morphology, from a disordered isotropic state to a highly ordered phase, quantitatively confirmed by a sharp increase in birefringence. Critically, we demonstrate the preservation of collagen’s native, periodic 66 nm d-banding, an essential motif for cell interaction. This structural refinement translates to a dramatic enhancement of mechanical performance, with the strongest filaments achieving an ultimate tensile strength of 0.64 GPa and a Young’s modulus of 8.9 GPa, values that surpass native tendon. Furthermore, the fabricated filaments are highly biocompatible and bio-instructive, promoting robust alignment and elongation of mesenchymal stem cells. This work provides a direct link between bioprocess parameters and the resulting hierarchical structure, offering a versatile platform for fabricating high performance, biomimetic materials for regenerative medicine.

## Introduction

The remarkable regenerative capabilities of human tissues have long fascinated civilisations. In the Greek tale of Prometheus, his liver would regenerate overnight to provide an eagle with eternal food, and torture Prometheus for eternity [1]. Beyond the intuitive grasp of regeneration that the myth offers, archaeological evidence does indeed show that by 3000 BCE humans engaged in primitive tissue grafting, where shells, gourds and silver/gold plates were used to reconstruct skulls [2, 3]. In fact, the oldest known surgical text, the Sushruta Samhita, from the 6th century BCE, documents detailed techniques for nasal reconstruction using forehead skin flaps, methods that not only restored form but also function [4]. Centuries later, in 1869, Jacques-Louis Reverdin pioneered the first successful skin autograft to treat burns and open wounds [5]. However, it was not until 1871, that the British surgeon George Pollock noted that autografts were superior to homografts, as he observed that when treating burn wounds on the same patient with autografts and homografts, skin treated with autologous grafts healed, whereas homografts would gradually “disappear” as the host rejected the transplant [6]. This superbly complex structural, mechanical, and biochemical microenvironment governs cellular behaviour and as a result, tissue functionality. By replicating these native conditions, tissue engineering strategies can overcome critical limitations such as immune rejection, mechanical mismatch, and poor integration with host tissues.

Recent advances in tissue engineering and regenerative medicine have demonstrated promising success in mimicking native human tissues. Hydrogels, for example, have been used as scaffolds to guide and organise encapsulated cells [7–9] with encouraging results. Among others, Niethammer et al. reported significant improvements on patients that received autologous chondrocyte hydrogel implants to treat knee joint cartilage defects [10]. Other approaches explored the use of decellularised tissue as templates for regeneration, where scaffolds maintain extracellular macromolecules such as collagen, elastin, fibronectin, and laminin, and provide tissue-specific cues for regeneration. For instance, Jiang et al. used decellularised placenta to create cardiac patches for myocardial repair after infarction [11], and Knox et al. first decellularised pulmonary artery grafts, re-endothelialised with host cells in a bioreactor and transplanted them [12].

More recently, bioprinting has emerged as a revolutionary technique for producing complex 3D tissue constructs. Breakthroughs include the development of a portable bioprinter pen for irregularly shaped wound [13], 3D-printed fully enclosed skin gloves for severely burned hands [14] and functional intestinal tubes, vascular junctions, and even a pumping heart model [15]. While these approaches provide significant control over both macro- and micro-architecture of engineered tissues; the choice of building material is equally critical for success biological integration. To create constructs that are not just structurally similar but truly functional, the ideal biomaterial should mimic the primary component of the natural tissues being replaced. Collagen undoubtably emerges as a prime candidate due to its biocompatibility, low immunogenicity, and ability to promote cell growth and attachment [15–17].

Collagen is an exquisite structural material. It represents 30% of the human body’s dry weight[18] and is able to deliver a wide range of mechanical profiles: it comprises most of the organic matter in skin, tendons, bones, and cartilage. The structural hierarchy inherent in collagen fibres, governed by intricate nanoarchitectural arrangements, plays a pivotal role in dictating the bulk mechanical properties of collagenous matrices [19–21]. Indeed, it is its fibrous nature that imbues it with a diverse array of properties, crucial for various physiological functions, including providing structural support to tissues like skin, cornea, cartilage, vasculature, and bones.

However, despite the impressive recent achievements, man-made collagen-based scaffolds consistently exhibit severely compromised mechanical properties that fundamentally limit their utility in load-bearing applications. In basic formulations, collagen hydrogels show Young’s moduli in the range of single digit kPa [22, 23], while gels with additional cross-linking stiffened only to a few tens of kPa [24–26]. In contrast, native connective tissues such as ligaments or tendons display stiffnesses of up to a few GPa and can hold forces up to several thousand Newtons before rupturing [27–30]. The mechanical strength of such tissues is greatly correlated with consistent directional organisation throughout their hierarchical structure, where collagen molecules assemble intro microfibrils, which form fibrils, which bundle into fibres, which in turn are packed into fascicles [31, 32].

Generating scaffolds that can simultaneously address nanoscale collagen organisation, microscale fascicle arrangement, and mesoscale architecture presents manufacturing challenges and has remained a barrier for current techniques [33]. Attempts to replicate this hierarchical structure for enhanced mechanical properties go back decades, with patents for collagen spinning going as far back as 1960 [34]. Techniques such as wet spinning or electrospinning offer good control over filament diameter and are highly scalable, unfortunately, single filaments often lack strength and flexibility and often require additional crosslinking using that introduce cytotoxicity [35–38]. Moreover, the use of harsh solvents and necessary high voltage often denatures collagen polymers [39, 40].

To overcome the challenge of creating tuneable, mechanically robust collagen scaffolds, we developed an accessible fibre pulling technique that integrates the precision of a commercial 3D bioprinter and leverages the physicochemical principles of molecular crowding. By extruding an acidic collagen solution into a hypertonic polyethylene glycol (PEG) coagulation bath, we aimed to induce rapid fibrillar assembly and alignment (Figure 1). In this study, we systematically characterise the resulting collagen filaments, detailing how these fabrication parameters directly influence filament diameter (Figure 2), surface morphology (Figure 3), structural organization (Figure 4), cellular interactions (Figure 5), and mechanical properties (Figure 6). This work establishes a scalable and versatile framework for fabricating high-performance collagenous materials, overcoming the critical limitations of conventional methods.

**Figure 1.**
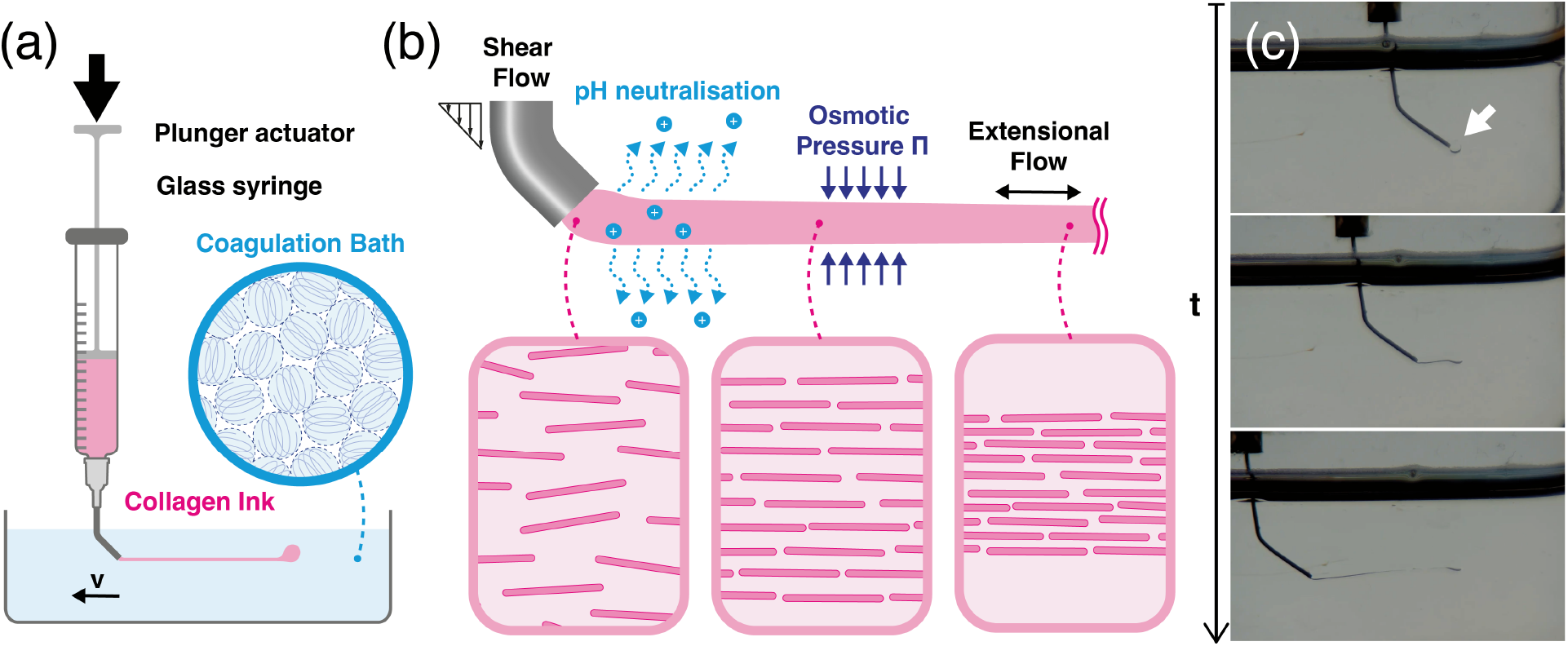
(a) Schematic representation of the fibre pulling method. Collagen is extruded into a polyethylene glycol (PEG) coagulation bath, where PEG polymers (blue) impose a colloidal trap that confines the collagen molecules (pink), preventing their diffusion. (b) Key physicochemical factors driving filament formation: (i) shear flow imposed by wall effects within the nozzle; (ii) neutralisation of the acidic collagen as protons diffuse into the surrounding bath (pH 7.4); (iii) osmotic pressure causing dehydration of the collagen phase; and (iv) extensional flow generated by the pulling motion. (c) Representative brightfield macrophotographs of the pulling process. An initial collagen blob is extruded to act as anchor (white arrowhead). After a 1 second settling period, the needle moves laterally at a constant speed, drawing a filament between the needle tip and the anchor point.

## Results

The formation of continuous collagen filaments in our system is driven by a combination of several physicochemical events that occur as collagen enters the polyethylene glycol (PEG) coagulation bath Figure 1(ab). To establish a predictable method for fabricating collagen filaments with tuneable characteristics, we first investigated the relationship between our primary process variables nozzle diameter and pulling speed and the resulting filament diameter. By systematically mapping the parameter space of eight commercial nozzle sizes and three distinct pulling speeds, we demonstrated precise control over filament dimensions, spanning nearly two orders of magnitude. For clarity, throughout the text we use the hierarchy **fibril > fibre > filament**, where fibrils are the smallest constituents, fibres are constructs of packed fibrils and filaments refer to bundled fibres.

### Filament size

We explored the parameter space encompassing eight sizes of commercial Luer lock blunt needles and three lateral speeds, and report a wide range of filament diameters, spanning from single to triple-digit microns. Filament diameters obtained using the pultrusion technique presented in Figure 2(b) demonstrated that filaments pulled from the most constrictive nozzles tended to be the thinnest, a trend consistently observed across pulling speeds. Furthermore, faster pulling speeds consistently yielded thinner filaments for all nozzle sizes. The smallest filaments within the parameter space were obtained using the smallest G34 nozzle (ID: 51 *µ*m) at the fastest pulling speed (10000 mm/min), resulting in filaments with diameters of 3.39 ± 1.3 *µ*m, representing the smallest published using off-the-shelf nozzles.

**Figure 2.**
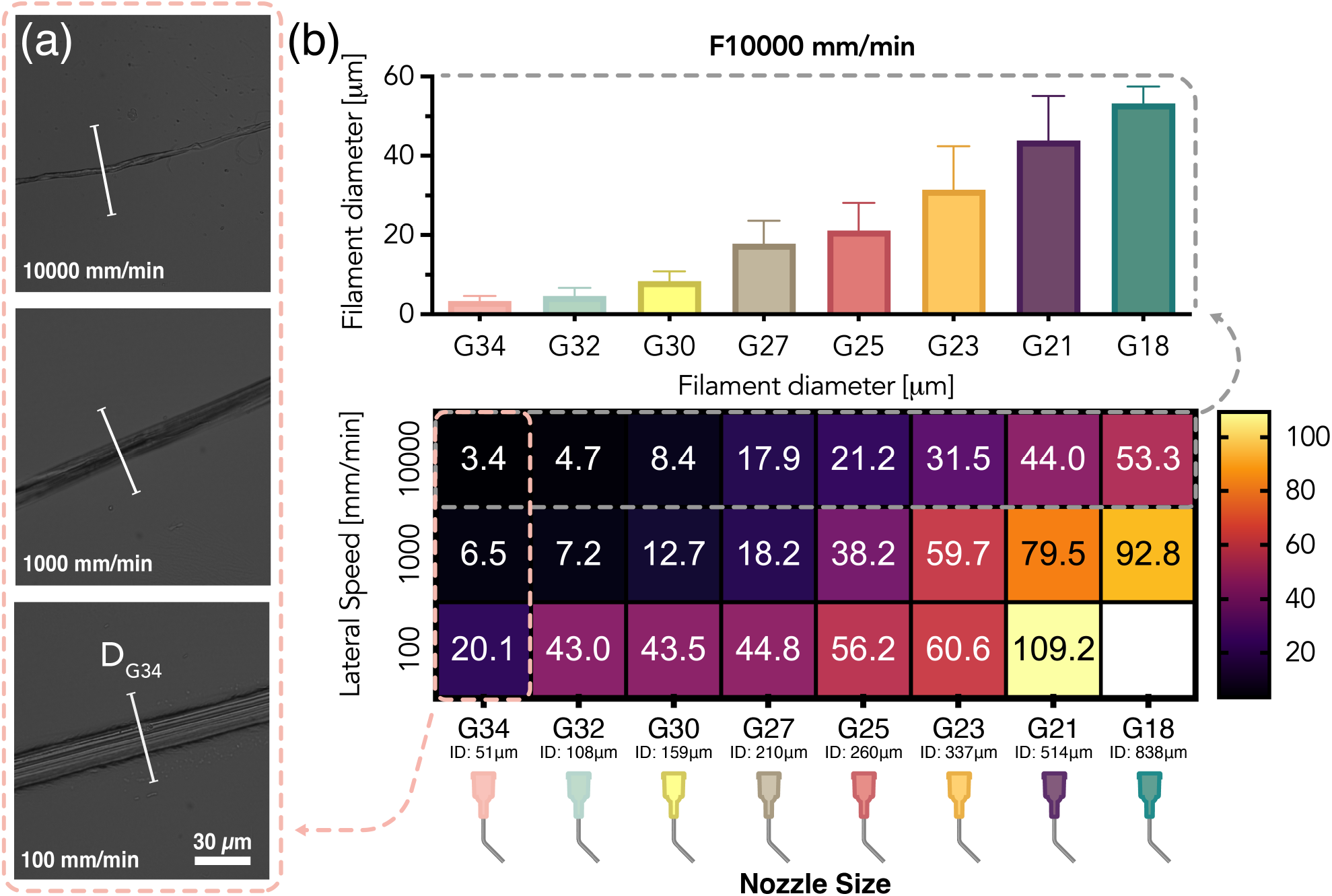
(a) Representative brightfield images of filaments pulled at the three tested feed rates using a G34 nozzle. To illustrate the relative size of filaments to the needle gauge, a white segment representing the nozzle size is included over each filament. (b) Influence of nozzle geometry and pulling speed on resulting filament diameter. The heatmap illustrates the average filament diameter obtained from permutations of 8 nozzle sizes (horizontal axis) and 3 lateral pulling speeds (vertical axis). The colour bar indicates the mean filament diameter in micrometres (µm). Each data point is the average of measurements from at least 10 independent filaments (n *≥* 10). Corresponding bar plots with statistical cross-comparisons are presented in S2.

Accelerating the pulling speed substantially reduced filament diameter, particularly in larger nozzles, while having a marginal effect on the smallest apertures, as illustrated in 2(b). Analysis of the effective circular cross-section across all filaments revealed tuneability of the filament/nozzle opening ratio from 15% to 0.2% (2(a), S1(b)). Interestingly, although the reduction in filament size with decreasing nozzle aperture may appear more pronounced for larger nozzles, the ratio of filament/aperture remained comparable.

To deconstruct the relative influence of our method variables, we analysed the correlation between fabrication parameters and the resulting filament. A multivariate Pearson correlation analysis unveiled that nozzle size was the dominant factor in controlling diameter, exhibiting a strong positive correlation with filament diameter (r=0.74). In contrast, feed rate had a smaller, negative correlation with filament diameter (r=-0.36). These results highlight the potential of a fabricating strategy using nozzle selection for coarse control and pulling speed for fine-tuning, offering precise control over the final filament dimensions.

Remarkably, even in the absence of fixed anchoring or specialised spinning setups, we successfully stretched filaments to sizes considerably smaller than the nozzle diameters. The notable exception was the permutation G18+F100 (largest nozzle at the slowest pulling speed), where filament formation was inhibited. Notably, we observed no evidence of necking in the collagen phase, hypothesizing that the substantial contact perimeter around the nozzle, combined with reduced drag resulting from slow pulling speed, impeded initial stretching of the collagen, causing it to be dragged along with the nozzle.

### Filament morphology

To comprehensively characterise the microfilaments, scanning electron microscopy (SEM) was employed to examine the topography of G18, G25, G30, and G34 filaments at pulling speeds of 100, 1000, and 10000 mm/min. Figure 3(c) presents a panel with representative images and directionality histograms. The SEM images unveiled significant variations in surface morphology, ranging from disordered isotropic appearance at the slowest settings to highly ordered grooves aligned along the pulling direction at the fastest settings.

**Figure 3.**
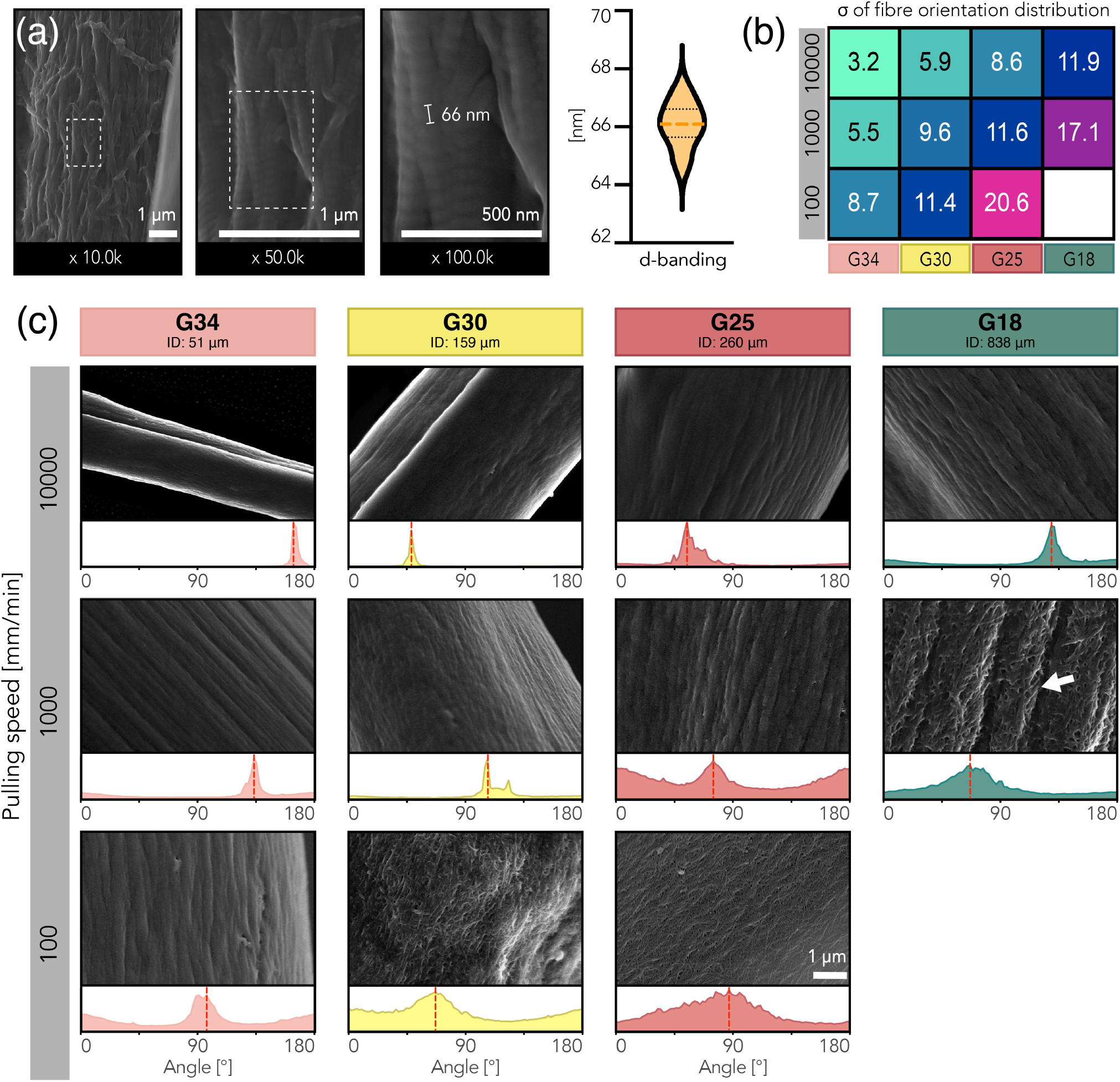
(a) Scanning electron micrographs of collagen fibrils within a filament at different magnification levels. Intact individual fibrils can be clearly observed across zoom levels. At 100,000x magnification, the distinctive collagen d-banding with 66-nm periodicity (n=18) is evident. The filament’s major axis is in the vertical direction.(b) Standard deviation of the global orientation obtained via Fourier spectrum analysis. Wider standard deviation is characteristic of the isotropic phase, whereas the more ordered phase displays narrower standard deviation. (c) Representative high magnification (10.000x) SEM images showing the varied fibrillar topographies of collagen filaments produced under different conditions. Arrowheads in the G18 F1000 sample highlight a uniquely interesting hierarchical arrangement with isotropic domains between aligned crevices. Surface morphologies are varied, ranging from wrinkled, corrugated appearance, in the bottom right of the panel, to a highly aligned, strained textile look for the top left filaments, suggesting a completed transition towards structural anisotropy. Nozzles are labelled with their gauge size, internal diameter, and colour code.

Across all filaments fabricated at a pulling speed of 100 mm/min (F100), minimal preferential direction was observed, indicative of an isotropic surface. Voids between fibrils were conspicuous, and the topography overall resembled the more usual collagen gel structure observed in previous work [23]. Fourier component analysis confirmed the absence of preferential order, with flat directionality histograms and comparably large dispersion for the F100 cases 3(b).

With increasing pulling speed, gaps and crevices became shallower, and improved alignment became evident. In the F1000 cases, crevices increasingly oriented along the filament’s main axis, with surfaces displaying a wrinkled appearance reminiscent of corrugated fabric, particularly evident in the G18 filament. The G18 F1000 filament exhibited a hierarchical arrangement with an isotropic-looking phase between aligned crevices, highlighted by arrows indicating rough surfaces between deep grooves. Progressing towards more constrictive nozzles, further topographical alignment was observed, with a transition towards a fully anisotropic phase becoming apparent, particularly in the G34 aperture.

Filaments fabricated at the fastest pulling speed (10000 mm/min) demonstrated improved directional order compared to all 100 mm/min cases. Surface morphology appeared smoother across all F10000 cases, further improving with decreasing filament diameter as grooves and voids between fibres became less apparent. This trend culminated with the G34-F10000 filament, exhibiting the smoothest, most densified, and homogeneous external layer. While alignment was comparable between G34-F1000 and F10000, the thinner F10000 filaments hinted at fibre compaction due to their almost halved diameter. This observation aligns with previous studies indicating the influence of fabrication parameters on the structural hierarchy and mechanical properties of fibrous materials [41, 42].

One of collagen I fibrils’ unique characteristics is the presence of d-banding; a repeating band pattern with a periodicity of 64-67 nm resulting from the quarter-staggered packing of collagen triple-helix molecules [43–46]. This periodic structure is conserved across different tissues and species [47, 48] and provides crucial topographical cues that control cellular behaviour such as adhesion, proliferation [49], contact guidance [45], and directional morphology and persistence [50]. High-magnification SEM images of our collagen type I filaments (Figure 3(a)) revealed the consistent presence of distinctive d-banding with a spacing of 66.07 nm (±0.78 nm), in line with the expected quarter-staggered molecular arrangement of collagen triple helices. This structural periodicity was observed uniformly across multiple filament samples (n=17), thus indicating the successful retention of the native fibrillar ultrastructure during fabrication.

### Structural organisation, alignment and order

By measuring the light retardance, polarized microscopy can provide a quantitative assessment of collagen alignment, structural anisotropy and thickness. Qualitatively, when inspecting a filament, bright areas correspond to strong retardance (Figure 4(a)). On the other hand, dimmer areas signal weaker retardance, suggesting lower collagen content, lack of consistent orientation, or both. To compare across conditions, the sample birefringence Δn is derived by dividing the retardance Γ by the optical length t (i.e. filament thickness).

**Figure 4.**
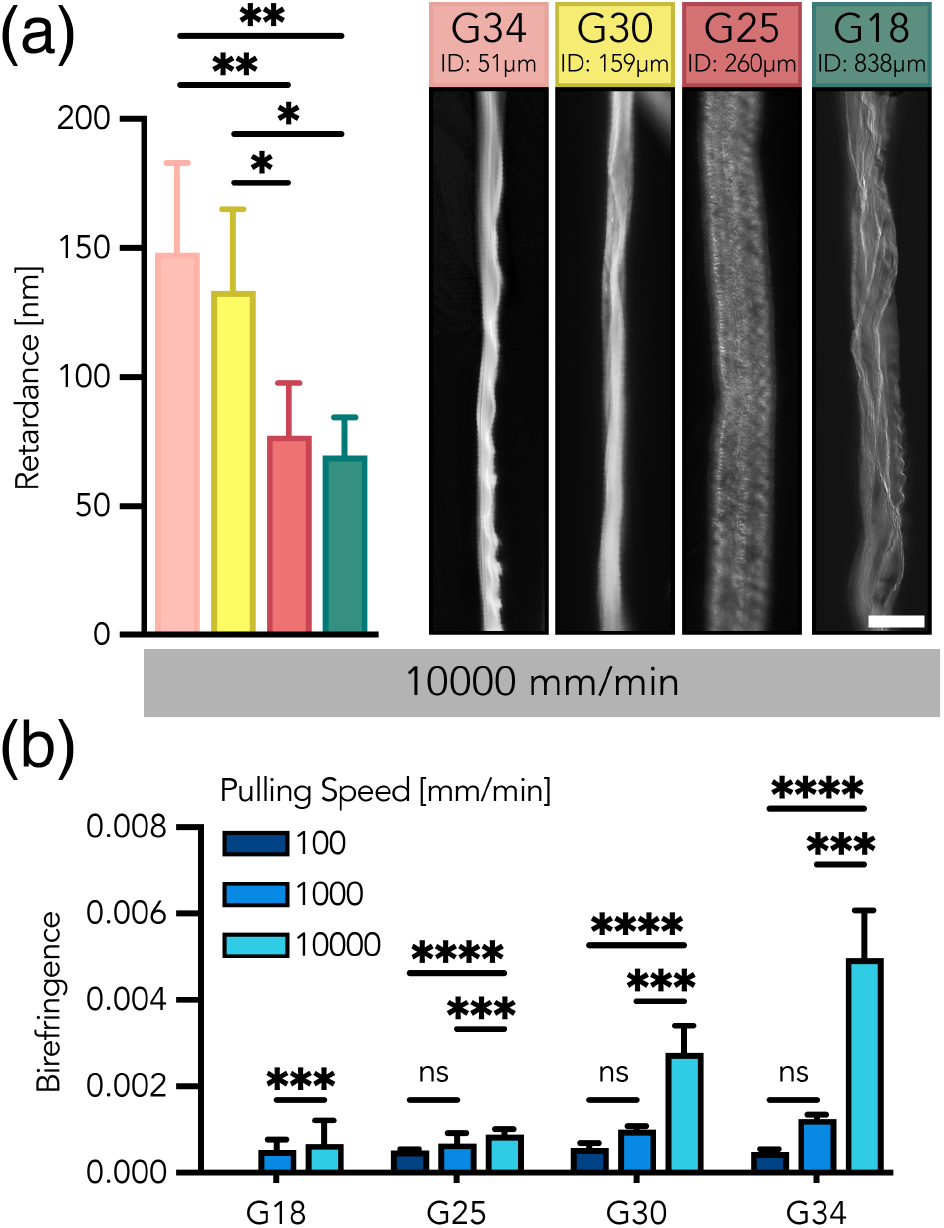
(a) Left. Light retardance of collagen filaments pulled at 10000 mm/min. Right. Representative polarised light microscopy images of F10k filaments. Intensity indicates the degree of light retardance (nm), where higher intensity indicates greater molecular compaction and alignment. Scale bar, 50*µ*m.(b) Quantitative analysis of filament birefringence as a function of nozzle size and pulling feed rate (100, 1000, and 10,000 mm/min). Birefringence was obtained by dividing each filament’s retardance by its diameter. Each data point represents the mean of at least 4 filaments (n *≥* 4), with measurements taken at 4 locations along each filament. Error bars indicate standard deviation. Asterisks (*) denote statistical significance (* p < 0.05, ** p < 0.01, *** p < 0.001,****p < 0.0001).

Filaments pulled at the lowest speed (100 mm/min) display weak birefringence, consistent with the largely unordered fibril arrangement observed in SEM images. Birefringence values are consistently low across all nozzles, suggesting little compaction and alignment (Figure 4(b)). At moderate pulling rates (1000 mm/min), birefringence shows non-significant increases for all 4 tested nozzles and remains low. At 10000 mm/min, filaments pulled using the more constrictive nozzles show a sharp increase in birefringence. The increased birefringence suggests a jump in molecular alignment and compaction, which qualitatively agrees with the topography of those filaments under SEM.

Overall, filaments show higher birefringence for higher pulling speeds across all nozzles. In addition, for the 1000 mm/min and 10000 mm/min cases, birefringence does increase progressively from the thickest nozzle condition (G18), to the more constrictive (G34).

### mMSCs alignment and morphology

To interrogate the influence of substrate dimensionality and curvature on cellular behaviour, we cultured murine mesenchymal stem cells (mMSCs) on thick and thin collagen filaments, using a collagen-coated glass-bottom plate as a 2D control. Here, we term “thick” as diameters significantly larger than the average MSC size (i.e. »20 µm), offering a gently curved surface, and “thin” as sub-cellular diameters with highly curved topography. Representative confocal images of mesenchymal stem cells at day 4 in all three culture conditions are shown in Figure 5(a). Cells in the control case (i.e. collagen coated petri dish) exhibit the usual spread for an isotropic substrate, showing polygonal morphologies with randomly oriented actin fibres. The absence of an underlying organised topography is confirmed by the broad distribution of cell orientation angles with no preferred orientation (Figure 5(b)). Cultured on a flat substrate, mMSCs display the lowest average cell aspect ratio (AR) of 2.75 ± 0.16 (Figure 5(c)) and the most circular nuclei (0.86) (Figure S4).

**Figure 5.**
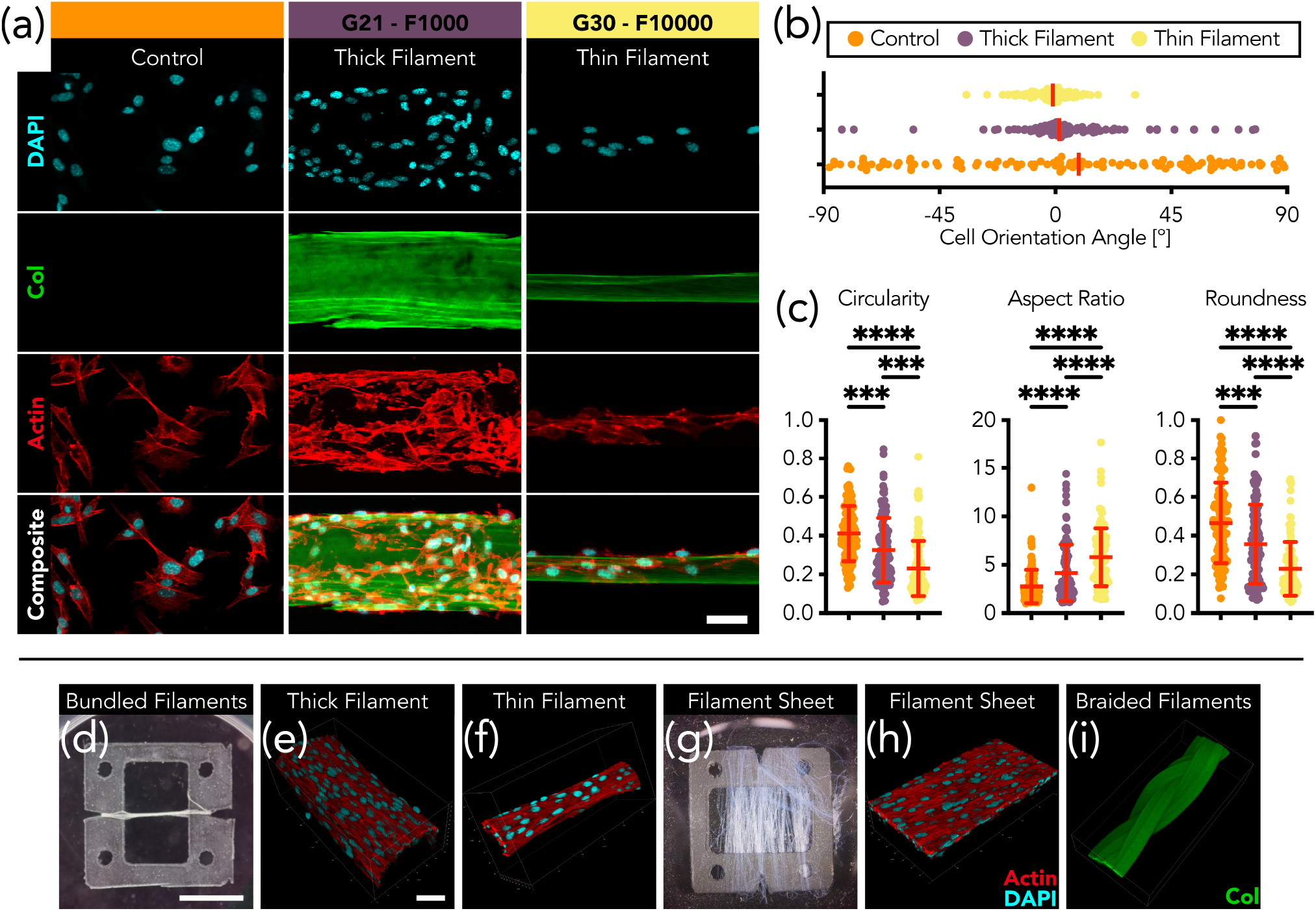
Morphological response and contact guidance of mouse mesenchymal stem cells (mMSCs) on collagen filaments. (a) Representative immunofluorescence confocal images of mMSCs after 4 days of culture on a 2D collagen-coated glass control and on pulled collagen filaments. Cells on the 2D control appear randomly oriented with a polygonal morphology, whereas cells on filaments align and stretch along the underlying filament axis. Staining: F-actin (red, via transfection), collagen (green, pre-stained with Alexa Fluor 488), and nuclei (cyan, Hoechst). Scale bar, 50 µm. (b) Quantification of cell body orientation relative to the filament’s major axis, demonstrating strong alignment on filaments compared to the random distribution on the 2D control. (c) Analysis of morphological descriptors. Cells on filaments become significantly less circular and more elongated (higher aspect ratio) compared to the 2D control, indicating a profound topographical response. Data is presented as mean ± standard deviation. Asterisks (*) denote statistical significance (***p < 0.001, ****p < 0.0001). (d) Closeup photograph of the custom laser-cut PVC stencil used for collecting and arranging filaments. The stencil is designed so that the corners contact the well walls of a 24 well plate. Triangular cutouts guide filament placement and help retain their original shape and length. Circular cutouts facilitate handling with tweezers. Scale bar, 60 mm. (e,f) 3D confocal reconstructions of confluent mMSCs enveloping (e) a thick filament and (f) a thin filaments after 7 days of culture. Scale bar, 50 µm. (g)A sheet-like lamina created by aligning multiple thin filaments along the edge of the collection stencil. (h)3D confocal reconstruction showing mMSCs forming a confluent monolayer across the filament lamina after 7 days. (i)3D confocal reconstruction of a braided, thread-like structure created by intertwining 3 individual thick filaments.

In contrast, cells seeded on both thick and thin filaments demonstrated a profound response to the topographical cues provided by the fibrous topography: mMSCs abandoned the random polygonal shape seen on the 2D control and instead, adopted a highly elongated, spindle-like morphology, aligning their major axis parallel to the underlying filament (Figure 5(b)). Contact guidance is particularly pronounced on thin filaments, where the difference between the average cell angle and the underlying filament is just 1.8 ± 0.87°.

As can be seen in Figure 5(a), cell circularity decreases from the control to the thick filament case (0.41 to 0.32), and from the thick to the thin filament case (0.32 to 0.23). Similarly, cellular aspect ratio increases monotonically as the available substrate for cell attachment narrows, peaking at 5.78 for cells on thin filaments. Interestingly, nuclei morphology appeared substantially different between the 2D control and filament cases, but not between filament conditions (Figure S4).

Cells remained highly viable (95% live cells) across conditions (Figure S4) and fully wrapped around filaments indicating that the collagen filaments offer favourable conditions for cellular growth and expansion. 3D reconstructions from confocal images in Figure 5(e) and (f), clearly shows how cells successfully attach and encapsulate both thick and thin filaments. Additionally, we also tested spooling filaments flat along the stencil to form a sheet (Figure 5(g)) and seed cells. By day 7, mMSCs had formed a confluent monolayer (Figure 5(h)) while retaining the alignment imposed by the underlying substrate.

### Mechanical characterization

The mechanical performance of fibrous constructs is governed by their organizational and structural hierarchy, which influences factors such as strength, modulus, extensibility, and toughness[51]. Understanding the relationship between structural features and mechanical behaviour is crucial for optimizing the design and performance of fibrous constructs. By systematically analysing the relationship between structural hierarchy and mechanical performance, we aim to elucidate key insights that can inform the development of advanced fibrous materials with enhanced mechanical properties.

Here, we investigate how filament thickness, compaction, and pulling speed affect the mechanical properties of fabricated fibrous materials. We characterised the strength, modulus and extensibility of filaments ranging from 3 to over 100 microns in diameter. Our results reveal novel insights into the relationship between structural hierarchy and mechanical behaviour, highlighting the importance of compaction in enhancing filament strength. Additionally, we observe a significant increase in Young’s modulus with decreasing filament diameter, underscoring the role of structural refinement in mechanical reinforcement.

A custom tensile testing rig with a 10-micronewton load cell was used to stretch individual filaments and examine their mechanical performance. A panel with representative stress-strain profiles is presented in Figure S5(a). All filaments were clamped at an initial length of 5mm and stretched at a constant rate of 0.1 mm/s. A single 29-micron filament (G25 – 10000) lifting a 2g weight without yielding is presented in Figure 6(b), demonstrating strength and handleability.

The most dexterous surgeons with the steadiest hands have tactile sensitivity of no less than 7.5mN [52, 53], so in applications where filaments have to be manipulated, maximum force values give an idea of how cautious the handling of these filaments must be: with G34 F100 being the most forgiving. The average maximum force is presented in Figure 6(d). Unsurprisingly, the isotropic and uncompacted G18 filaments underperformed all other cases. Interestingly, even though the average diameter of G18-10000 and G25-100 filaments are comparable (53.3 *µ*m and 50.4 *µ*m respectively), the rupture force was an order of magnitude superior in the G25 case despite larger variance in directionality, suggesting that compaction and not just directional order are necessary. A similar instance can be observed between G30-10000 (8.4 *µ*m) and G34-1000 (6.5*µ*m) where the thinnest filament was able to endure a larger force despite comparable directional order.

**Figure 6.**
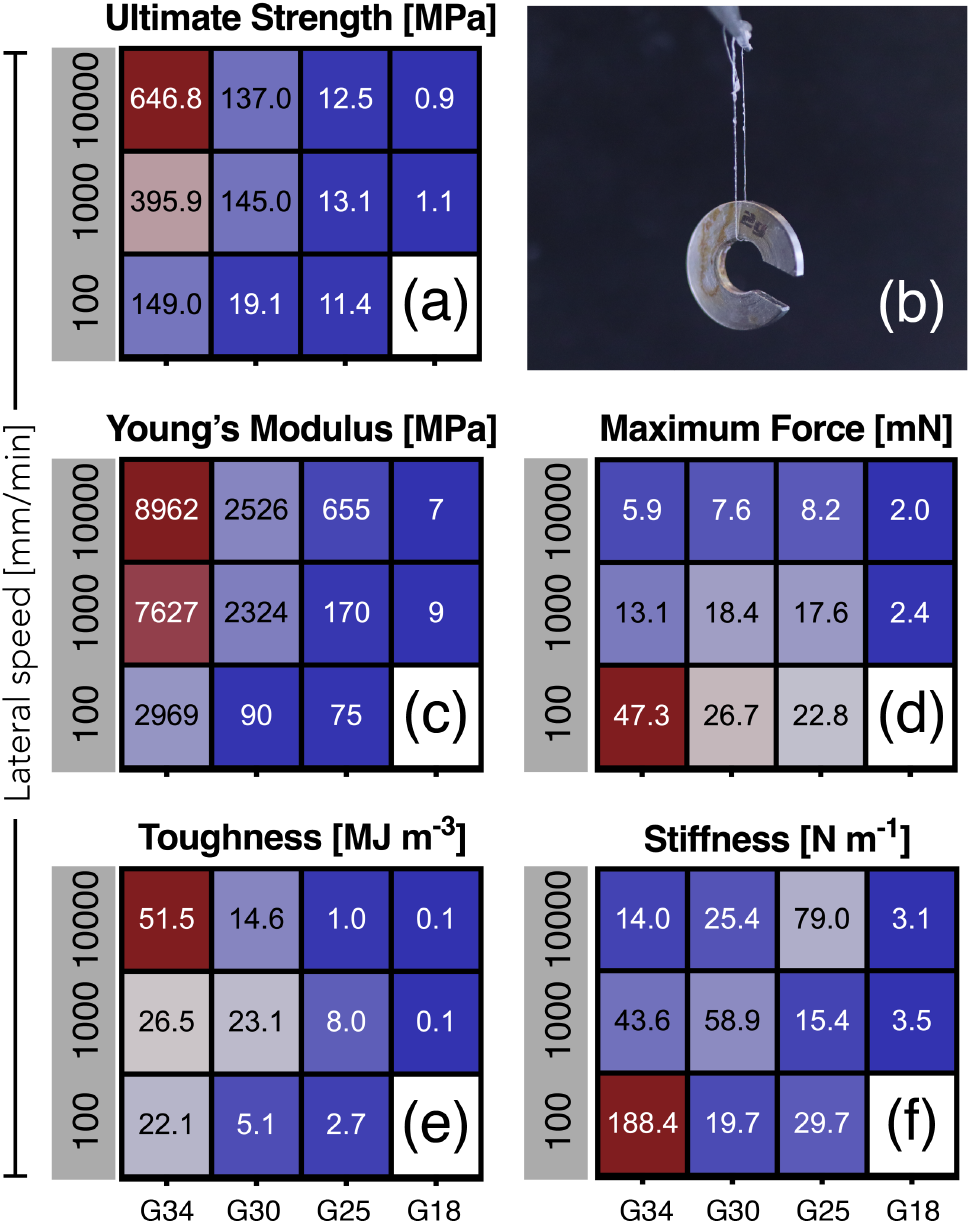
Comparison of mechanical characteristics of single fibres: (a) ultimate tensile strength, (c) Young’s modulus, (d) rupture force, (e) energy at rupture, and (f) stiffness. Each data point is the average of measurements from at least 5 independent filaments (n *≥* 5). (b) A single collagen filament (D = 29 *µ*m) lifting a 2g weight demonstrating handleability and strength. Grouped bar plots with cross comparisons for each of the mechanical metrics can be found in Figure S7.

Interestingly, while thicker filaments initially displayed greater strength in millinewton terms, normalisation to their initial cross-sectional area (engineering stress) revealed the inverse trend. Filaments pulled through more constrictive apertures, such as G30 and G34, exhibited increased ultimate tensile strength (UTS) with higher pulling rates. For instance, UTS values for G30 and G34 increased from 19 MPa to 137 MPa and from 149 MPa to 648 MPa, respectively (6(a)). Similarly, Young’s modulus showed a substantial increase with decreasing filament diameter (Figure 6(c), S5(b)), indicating enhanced stiffness with structural refinement. Our 3.4 *µ*m filaments, pulled at 10000 mm/min from a G34 nozzle exhibited superior mechanical performance compared to any previously reported collagenous constructs, with 8.9 GPa elastic modulus and 0.64 GPa UTS.

## Discussion

Collagen is an impressively versatile material, found in many body structures with a wide range of structural characteristics. In line with collagen’s physiological versatility, we used rat tail collagen I acidic solution, which in the precursor tendon form has a modulus of 498 MPa[54, 55], to fabricate constructs ranging from much softer to much stronger than the source tissue.

Collagen concentration in the dope solution is known to affect fibril assembly and spacing: the average distance between collagen molecules decreases as the volume fraction of collagen in the solution increases [43, 56, 57], while the fibre diameter and storage modulus increase with increasing concentration [58–64]. As a result, the specific strength of fibres increases as the precursor solution gets concentrated [65]. In both studies, PEG 20000 is used as the molecular crowding agent, however there is a wide difference between the concentration used in the “flow-focusing buffer” (10% w/v) and our “coagulation bath” (49% w/v). In polymer solutions, the osmotic pressure increases with concentration; linearly in the dilute regime and with a power of 5/4 in the semidilute regime [66] [67–71]. As a consequence of the high osmolarity around the collagen phase, the spacing between collagen molecules decreases [72, 73], resulting in thinner, more compacted fibrils with increased stiffness and indentation modulus [74].

More importantly, the effects of macromolecular crowding are marginal until the colloidal solution reaches its overlap concentration [75], which for PEG 20000 can be easily calculated as c* = 24 mg/ml [66, 67] [76–79]. Concentrations greater than c*, promote phase separation, confine the collagen within a “colloidal cage”, promote alignment and boost nucleation rate [66, 75, 80]. Binding minimises the volume from which collagen molecules are excluded, so the “excluded volume” effectively creates an attractive force that actively pushes collagen molecules, thus promoting initial assembly and further structural compaction of newly formed fibrils together [41, 78, 79, 81].

The selection of coagulant and concentration are critical considerations in the design of coagulation baths. They are, however, not sufficient, as illustrated by the striking differences between Haynl et al.[16] and Lu et al.[82] constructs. Both studies used the same PEG20000 concentration (10% w/v), however, Haynl et al.’s impressive megapascal-strong fibres are several orders of magnitude stronger than Lu et al.’s non-crosslinked fibres (E: 9.32 kPa, UTS: 3.2 MPa) and crosslinked fibres (E: 97.02 kPa). Other studies such as Yaari et al.[83] used a coagulation buffer of PBS and salts without PEG, and instead treated their wet-spun collagen fibres with glutaraldehyde (GTA) for additional cross-linking (UTS: 378MPa E: 3.5GPa). In contrast, Koeck et al.[44] coagulated their collagen I fibres in acetone, resulting in constructs with 240MPa strength.

This study presents a highly accessible and tuneable method for fabricating high-performance collagen filaments by combining the precision, convenience and accessibility of an off-the-shelf bioprinter with the physicochemical principles of molecular crowding. By systematically controlling just two parameters-pulling speed and nozzle diameter we produced un-crosslinked, biocompatible filaments with mechanical properties that span several orders of magnitude, in some cases matching or surpassing those of native tendons. This work offers a powerful yet gentle method to engineering high aspect ratio collagenous biomaterials, avoiding the harsh chemical treatments and solvents that often compromise biocompatibility in traditional fabrication methods. This simple yet effective scheme offers significant advantages over other methods that may require complex microfluidic setups or chemical modifications to achieve a similar range of properties. Conventional wet spinning, for instance, often demands post-fabrication chemical crosslinking, to achieve comparable mechanical strength, which can introduce cytotoxicity and alter fibre morphology [40, 84, 85].

Our proposed approach also overcomes the primary drawbacks of electrospinning. Similarly, in electrospinning, the combination of harsh solvents and high voltage fields often denatures collagen’s native triple-helical structure, turning it into gelatin-like random coils [39, 86–88]. As a result, the native 67 nm D-banding, essential for cell recognition, is disrupted, resulting in diminished mechanical integrity and loss of essential bioactive signals [40, 87, 89]. By contrast, our method, by using an aqueous-based crowding environment, successfully preserves this native ultrastructure, creating filaments that are both mechanically robust and biologically active without the need for blending collagen with other polymers or subsequent chemical treatment.

We recruit and combine the effects of shear alignment, extensional flow, osmotic pressure, and molecular crowding to adjust collagen’s morphology and alignment. The PEG molecular crowding environment enhances collagen fibrillogenesis through excluded volume effects, accelerating nucleation and promoting fibril alignment. By capitalising on the quasi-instantaneous gelation facilitated by the coagulation bath, we show that the interplay between nozzle size and pulling speed offers a robust framework for controlling filament diameter, which in turn dictates its structural and mechanical properties.

Our thick filaments, with single-digit modulus and sub-MPa strength, stand out as considerably softer than most filaments reported in the literature where any form of molecular crowding or coagulation is involved [44, 82, 90]. Only Lu et al.[82]’s 100 *µ*m non cross-linked filaments show a similar-low- strength of 0.5MPa. Wet filaments produced by Koeck et al.[44] through a G18 nozzle (100 *µ*m), equivalent to our G18-F1000, are 10 times higher strength (13MPa) and 4 times stronger in modulus (42MPa). On the other hand, our thinnest filaments, with single-digit-micron diameters, are within the strongest reported in the literature. Our 3.4 *µ*m filaments (i.e. G34-F10k) are comparable to the ones Haynl et al.[16] produced from 5mg/ml collagen I, using a hybrid device that combined wet spinning and flow focusing. Interestingly, even though the diameter of our filaments (3.4 *µ*m) and their microfibres (3.7 *µ*m) are comparable, the resulting mechanical characteristics diverge slightly: 383MPa and 4138MPa tensile strength and modulus for their microfibres; compared to the 646MPa strength and 8962MPa modulus we found for our filaments.

Polarised light microscopy (PLM) has been a standard technique used to inspect molecular order and feature alignment since the XIXth century [91]. Collagen exhibits intrinsic birefringence due to its asymmetrical triple-helical structure, and form birefringence, which arises from the geometric ordering of molecules in space [92, 93]. When collagen molecules are coherently aligned and densely packed, their intrinsic birefringence add up. As fibrils align, their intrinsic birefringence are superimposed, resulting in higher retardance-proportional to the degree of molecular organisation within collagen filaments-. Conversely, a randomly oriented network exhibits low retardance as the optical contributions cancel out. The measured optical retardance serves as a direct proxy for both molecular alignment and packing density [94–96]. Therefore, the sharp increase in birefringence observed with smaller nozzles and faster pulling speeds quantitatively confirms the formation of densely packed, highly anisotropic structures [96–99].

In agreement with previous work, our results highlight the pivotal role of structural refinement in enhancing the mechanical performance of collagenous filaments [44, 83, 90]: in highly birefringent filaments, constituent collagen molecules are aligned along the filament’s long axis, thus, as the filament is loaded in that direction, the alignment and packing allow for the efficient distribution of stress, resulting in higher Young’s modulus and ultimate tensile strength [100–103]. In contrast, isotropic, low-birefringence filaments must first deform as its fibrils re-align, before the molecular backbones bear the load, hence yielding lower stiffness and strength [104].

The mechanical performance of our filaments is matched by their biological relevance, underscored by the preservation of collagen’s native ultrastructure. High-magnification SEM imaging confirmed the presence of the characteristically periodic 66 nm D-banding across our filaments, which provides critical topographical cues for cell-matrix interactions through contact guidance [45, 49, 105]. While cells are known to sense and respond to substrates with periodicity in the range 35-250 nm [106–111], collagen I d-banding in native human tissues sits in a much narrower range of 64-67nm [47, 48, 112]. Crucially, our fabrication method faithfully recreates the specific 66 nm bio-signature found in native tissues. The robust alignment and highly tensional phenotype exhibited by mMSCs cultured on our filaments are direct manifestations of this high-fidelity guidance. Nevertheless, the significance of achieving physiological spacing on our filaments extends beyond cell alignment or elongation. By maintaining this bio-instructive signal, our filaments provide a truly biomimetic scaffold and could set the groundwork for guiding critical complex cellular processes required to engineer truly functional, load-bearing tissues capabilities often lost in harsher fabrication methods.

## Conclusion

Our study elucidates the powerful influence of structural hierarchy on the mechanical and biological performance of fibrous biomaterials. We have developed and characterised a scalable and accessible method that leverages molecular crowding and 3D-bioprinting to fabricate collagen filaments with precisely tailored features. We demonstrated that adjustments on just two mechanical inputs-nozzle size and pulling speed can be leveraged to tune structural refinement and compaction, significantly enhancing filament strength and stiffness while preserving native biocompatible motifs. These findings contribute to a deeper understanding of the relationship between structural features and mechanical behaviour in engineered fibrous constructs and offer a valuable platform for designing advanced, cell-instructive materials for regenerative medicine. Further investigations into the underlying mechanisms governing the relationship between structural hierarchy and mechanical behaviour are warranted to advance the design and optimisation of fibrous materials for medical applications.

## Methods

### Collagen ink

Acetic-acid-solubilised type I rat tail collagen (Corning, 354249) was used in the experiments. The 10.1 mg/ml collagen solution was loaded into a 2.5 ml Hamilton glass syringe, which was then mounted onto a custom gantry carriage. This assembly was specifically designed to incorporate a Peltier cooler and heatsink to regulate the temperature of the collagen ink within the syringe, maintaining it at 5°C throughout the experimental procedure.

### Coagulation Bath

All experiments were carried out using Poly(ethylene glycol) with average *M*_*w*_ 20,000 Da (Sigma-Aldrich, 25322-68-3), hereafter PEG-20k, as the crowding agent. The stock coagulation bath was prepared by mixing powdered PEG-20k in phosphate buffered saline (PBS 1X, pH 7.4) (Gibco, 10010-023) to a final mass-per-volume concentration of 480 mg/ml. This solution is shelf-stable and can be stored in a sealed container at room temperature.

### Fabrication of collagen filaments

Off the shelf 18, 21, 23, 25, 27, 30, 32 and 34-gauge nozzles were used to deliver collagen. The nozzles were gently bent 45 degrees around the middle to reduce damage of the collagen phase as it left the nozzle. A commercial open-source syringe bioprinter (Lultzbot Bio) with the aforementioned Peltier cooler was used to fabricate the filaments. Custom G-code of the form *G0 X60 F1000* was used to pattern 60mm filaments at three different feed rates: 100, 1000 and 10000 mm/min. To pull collagen filaments, anchoring blobs were extruded and allowed 1 second to gel before the pulling motion started 1(c). The collagen blobs were generated using a command of the form *G1 E0*.*1 F10*, which resulted in a blob size of 4.17 *µ*l S1(a). The apparent arbitrariness of the anchor volume was the result of the minimum movement of the extruder axis in the LulzBot Bio (0.1 mm). To facilitate handling and observation, an additional anchoring blob was extruded at the end of each filament. Filaments remained in the coagulation bath for at least 2 minutes before they were collected with laser-cut stencils, carefully lifted and washed in a PBS bath to remove excess PEG-20K.

### Cell culture and maintenance

Bone marrow-derived mesenchymal stem cells (mMSCs; D1, ATCC, CRL-12424, Manassas, VA, USA), previously transfected with LifeAct-RFP, were maintained under standard culture conditions at 37^°^C and 5% CO_2_. All experiments used cells at passage number 6 or lower to ensure consistency and minimise potential phenotypic drift. The culture medium consisted of Dulbecco’s Modified Eagle Medium (DMEM; ThermoFisher Scientific, NY, USA, Cat. #11965-092) supplemented with 10% fetal bovine serum (FBS; Corning, MA, USA, Cat. #35-015-CV) and 1% (w/v) penicillin/streptomycin (P/S; ThermoFisher Scientific, USA, Cat. #15140122), referred to as complete DMEM medium. Tenogenic media was prepared by supplementing DMEM complete medium with 50 µg/mL ascorbic acid (AA; Sigma, St. Louis, MO, USA, Cat. #95209-50G), 50 ng/mL GDF-5 (R&D, MN, USA, Cat. #853-G5), 100ng/mL CTGF (Acro, DE, USA, Cat. #CTF-M52Hc) and 10 ng/mL TGF-β3 (Stemcell Technology, USA, Cat. #78131). Differentiation protocols were initiated one day post-seeding by replacing the complete medium with appropriate differentiation medium, designated as Day 0. Media was replenished every other day in the first 7 days and every day in the subsequent 7 days. Corresponding control samples maintained in complete medium were collected at matching time points for comparative analysis.

### Immunofluorescent staining

For immunofluorescent analysis, cells were fixed with 4% paraformaldehyde (Santa Cruz, TX, USA, Cat. #281692) for 15 minutes at room temperature and permeabilised with 0.3% Triton-X100 (Sigma, MA, USA, Cat. #9036-19-5) in PBS for 15 minutes. Cells were then blocked with 1% (w/v) BSA (Sigma, St. Louis, MO, USA, Cat. #A9647-100G) in PBS for 1 hour at room temperature. The tenomodulin (TNMD) antibody was diluted 1:100 in 1% BSA (Invitrogen, USA, Cat. #PA5-112767) and incubated at 4°C overnight. Alexa 488 (Invitrogen, USA, Cat. #A27034) and Hoechst (Invitrogen, USA, Cat. #H3570) were used for secondary staining at 1:500 and 1:2000 dilution respectively. Validation of the primary antibodies are provided in the manufacturers’ websites.

### Confocal microscopy

A Leica DMi8 SP8 scanning laser confocal microscope and the software Leica Application Suite X (LAS X, v. 3.5.7.23225) were used to image all brightfield and fluorescent samples. Samples were maintained in glass-bottom dishes during image acquisition. A multi-channel line sequence scanning protocol was used to prevent spectral bleed-through. Collagen filaments were imaged using 10x and 20x objectives (Wetzlar, Germany). mMSCs were imaged at higher resolution using the 20x objective and taking z-stacks with 5*µ*m z steps. For 3D visualisation, z-stacks were acquired using either a 20x objective or 40x oil immersion with 1*µ*m steps through the entire volume of cells and constructs. For quantitative analysis, maximum intensity projections of the z-stacks were generated. Cell outlines were traced manually using an iPad Pro and shape descriptors were extracted using the open-source software Fiji (ImageJ). Cellpose [113] was used to segment cell nuclei and Fiji to extract shape descriptors.

### Polarised light microscopy

The optical anisotropy collagen filaments was quantified using a Nikon Eclipse TE200 inverted microscope equipped with a rotating circular polarizer, an analyser, and a liquid crystal compensator (LC-PolScope, Abrio). Filaments were imaged using a 20x/0.4 NA objective (0.3µm/pixel), fitting the entire width of each filament in a single field of view. The average optical retardance (Γ, in nm) was measured across 4 different locations on each filament to ensure representative sampling. Birefringence (Δ*n*) was computed according to the formula: Δ*n* = Γ*/t*, where Γ is the measured retardance, and *t* the optical path length, in this case the filament’s diameter. Filament diameters were measured from corresponding brightfield images taken on the same system.

### Scanning Electron Microscopy

Collagen filaments used to assess directionality were first removed from the coagulation bath and thoroughly rinsed in phosphate-buffered saline (PBS). Subsequently, the samples underwent a 5-minute soaking period in diH2O before being mounted onto glass substrates and air-dried overnight. Collagen filaments used to inspect d-band spacing were fixed in 2.5% glutaraldehyde solution (Thermo Fisher Scientific) and maintained at room temperature for a minimum of 16 hours to ensure structural preservation and stabilization. Following the fixation period, samples were washed with deionized water through a series of four sequential washes: three 10-minute washes followed by one extended 30-minute wash to remove residual fixative. Complete dehydration was achieved through a series of graded ethanol washes to prevent collapse and maintain structural integrity: 50% ethanol (two sequential treatments), 70% ethanol (two sequential treatments), 90% ethanol (two sequential treatments), 95% ethanol (two sequential treatments), and 100% ethanol (three sequential treatments). Each dehydration step was performed for 15 minutes to ensure uniform solvent exchange throughout the sample. To limit surface tension artifacts, samples were further dried through a sequence of hexamethyldisilazane (HMDS) graded washes: 2:1 ratio of ethanol to HMDS for 15 minutes, 1:1 ratio treatment for 15 minutes, 1:2 ratio treatment for 15 minutes, and finally three sequential 15-minute treatments in 100% HMDS. Samples were mounted onto carbon tape-covered stubs to prevent deformation during solvent evaporation and were then allowed to dry overnight in a chemical fume hood under ambient conditions. The dried samples were coated with a 5 nm layer of iridium using in a high-vacuum sputter coater for 60 seconds. The morphology of the filaments was characterized using a Hitachi SU-70 scanning electron microscope (Yale Institute for Nanoscience and Quantum Engineering) at a 5 kV acceleration voltage. Images were exported as high resolution (1280×960 pixels) TIFF files to prevent data loss from compression and preserve the fine details for analysis. The alignment of surface features was quantitatively analysed using the Directionality plugin in Fiji (ImageJ2, v2.16) in x10.0k SEM images. This specific magnification was deliberately chosen as it provided sufficient resolution to capture topographical features at a length scale relevant to cell-matrix interactions, while simultaneously maintaining a field of view large enough to observe the overall supramolecular organisation of fibres/fibrils. The Directionality plugin computes a directionality histogram based on the 2D Fast Fourier Transform (FFT) power spectrum of the image. To ensure the analysis was restricted to relevant features, masks were created for each image to isolate the in-focus filament surface and exclude the background (Central column in Figure S3). The analysis was performed with 90 angular bins. Instead of using the plugin’s default Gaussian fit dispersion output, the raw directionality histograms were exported for each sample. The standard deviation of each orientation distribution was then calculated to serve as a quantitative metric for alignment dispersion. A lower standard deviation indicates a higher degree of fibril alignment (anisotropy), while a higher value reflects a more random, disordered orientation (isotropy). The final dispersion value reported for each condition represents the average standard deviation from at least four independent filaments (n *≥* 4).

### Mechanical characterization

For each filament, the cross-sectional diameter was determined by averaging measurements in at least four different locations. At least six filaments were measured per condition. The inspect the surface topography, the ImageJ plugin Directionality was used. The reported dispersion was computed as the average of the standard deviation of the gaussian across samples. Filaments with a length of 60 mm were prepared for mechanical testing according to the standard ASTM C1557-20. Tissue adhesive (Liquivet, VVG3SC) was utilised to affix the filaments to a stencil. Once the glue had fully polymerized, the stencil containing the filament was mounted onto a custom tensile testing rig. To ensure accurate measurements and maintain filament integrity, the top jaw was locked first and lowered to a position 5 mm from the bottom jaw before securing the bottom jaw and cutting the sides of the stencil. Force readout was continuously monitored throughout the process to confirm that no force was applied to the filament. Tensile tests were conducted at a loading rate of 0.1 mm/s until rupture occurred. All tests were performed at a temperature of 21°C and a relative humidity of 90%. A detailed diagram describing the data processing pipeline is included in Figure S6. The raw Force-time profiles were converted to force-displacement using the known stretch rate. From the force-displacement data, we derived several key mechanical metrics. The Stiffness (N/m) was calculated from the initial linear slope of the force-displacement curve (below 2% displacement). The engineering stress-strain curve was then generated by normalising force by the initial cross-sectional area (*A*_0_) and displacement by the initial gauge length (*L*_0_). From that curve, we determined the Young’s Modulus (E) from the slope of the linear fit in the 0-2% strain region. We extracted the Ultimate Tensile Strength (UTS) from the peak stress, and the Tensile Toughness (*U*_*T*_) from the integrated area under the curve. At least five independent filament were tested (n *≥* 5) per condition.

### Statistical analysis

All statistical analyses were conducted using GraphPad Prism 10.4.2 software. Bar plots depict mean values with error bars representing the standard deviation (SD). Sample sizes and p values are detailed in the figure or figure legends. A significance threshold of p < 0.05 was used, with comparisons yielding no significant difference denoted as “ns”. For p values greater than 0.0001, they are explicitly stated either within the figures or in the accompanying legends.

## Notes

### Competing Interest Statement

The authors have declared no competing interest.

### Funding

- National Institutes of Health: R35GM142875

