## Supplementary Information for "Development of Collagenous Filaments with Tuneable Mechanical Properties Using a 3D Bioprinter and Molecular Crowding"

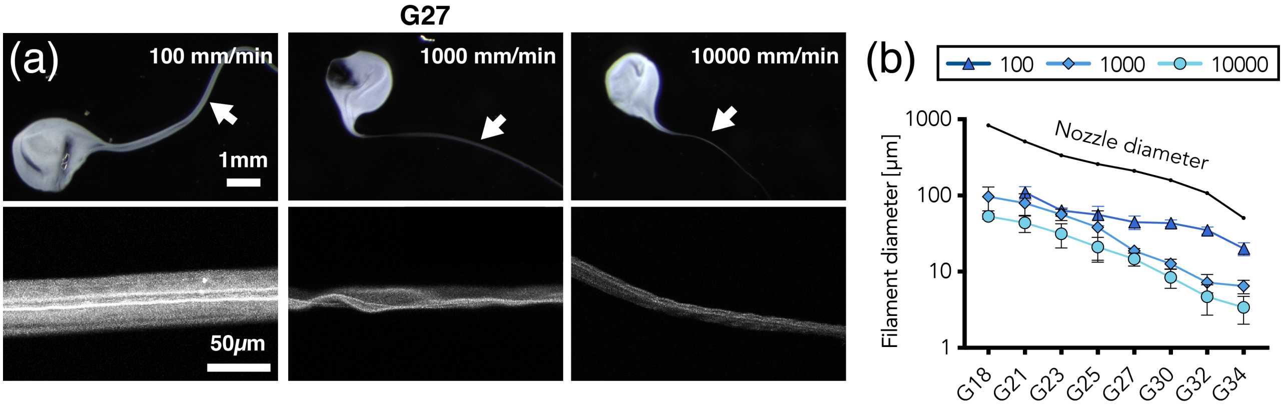


**Figure S1.** (a) Top row: Representative darkfield images of the blob and filaments constructs. Bottom row: Representative confocal images of the pulled filaments. (b) Filament diameter evolution for the three tested lateral speeds: 100, 1000 and 10000 mm/min across nozzle diameters.

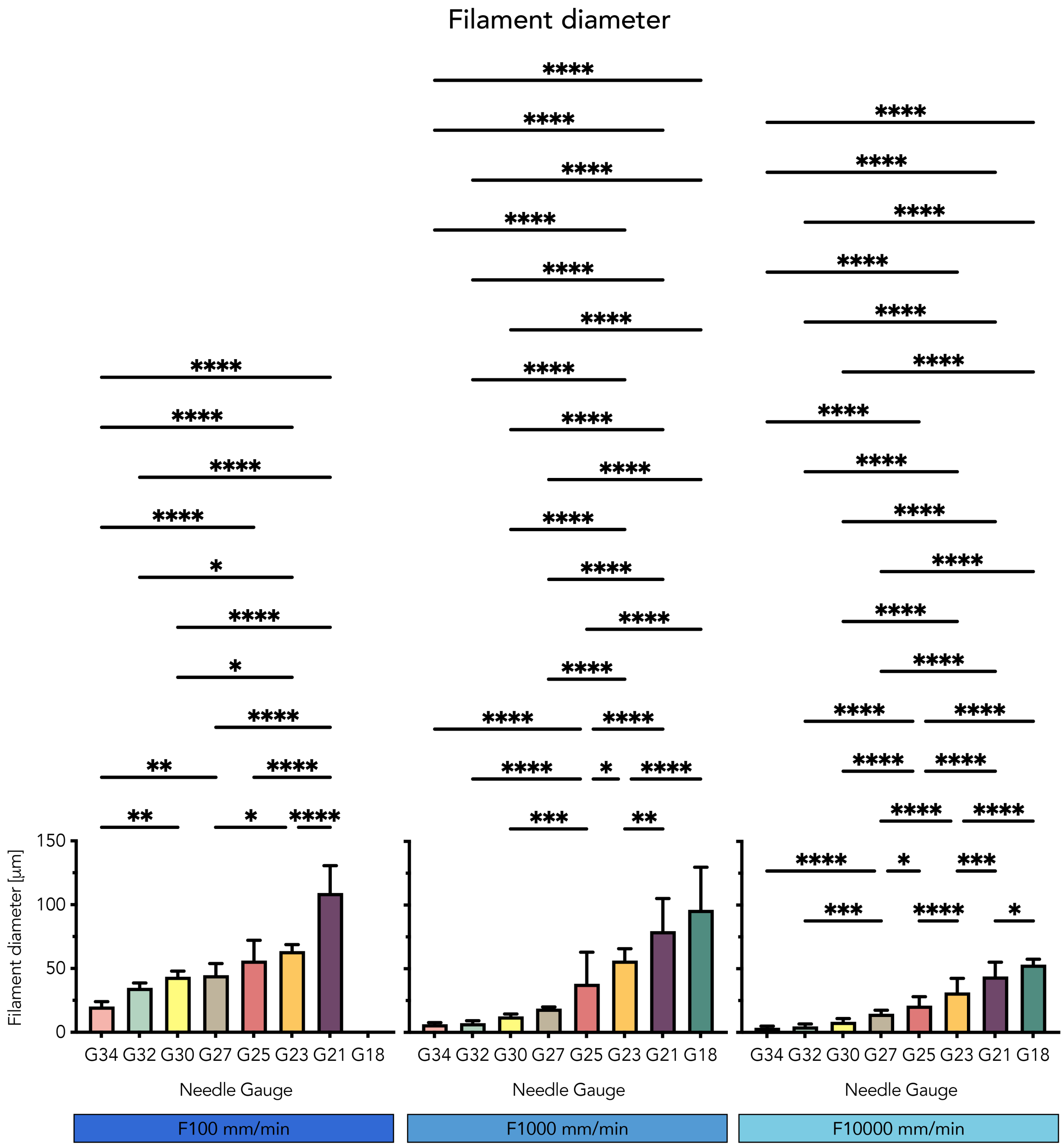


**Figure S2.** Influence of nozzle gauge on filament diameter at different pulling speeds: 100 mm/min (left), 1000 mm/min (center), and 10000 mm/min (right). Each bar represents the mean filament diameter (in µm). Error bars denote the standard deviation (SD) from at least 10 independent filament measurements ($n \geq 10$). Bars are color-coded to match the Luer lock nozzle colours. Horizontal lines indicate statistically significant differences between nozzle gauges, as determined by a one-way ANOVA. Asterisks (*) denote statistical significance (* $p < 0.05$, ** $p < 0.01$, *** $p < 0.001$, ****$p < 0.0001$).

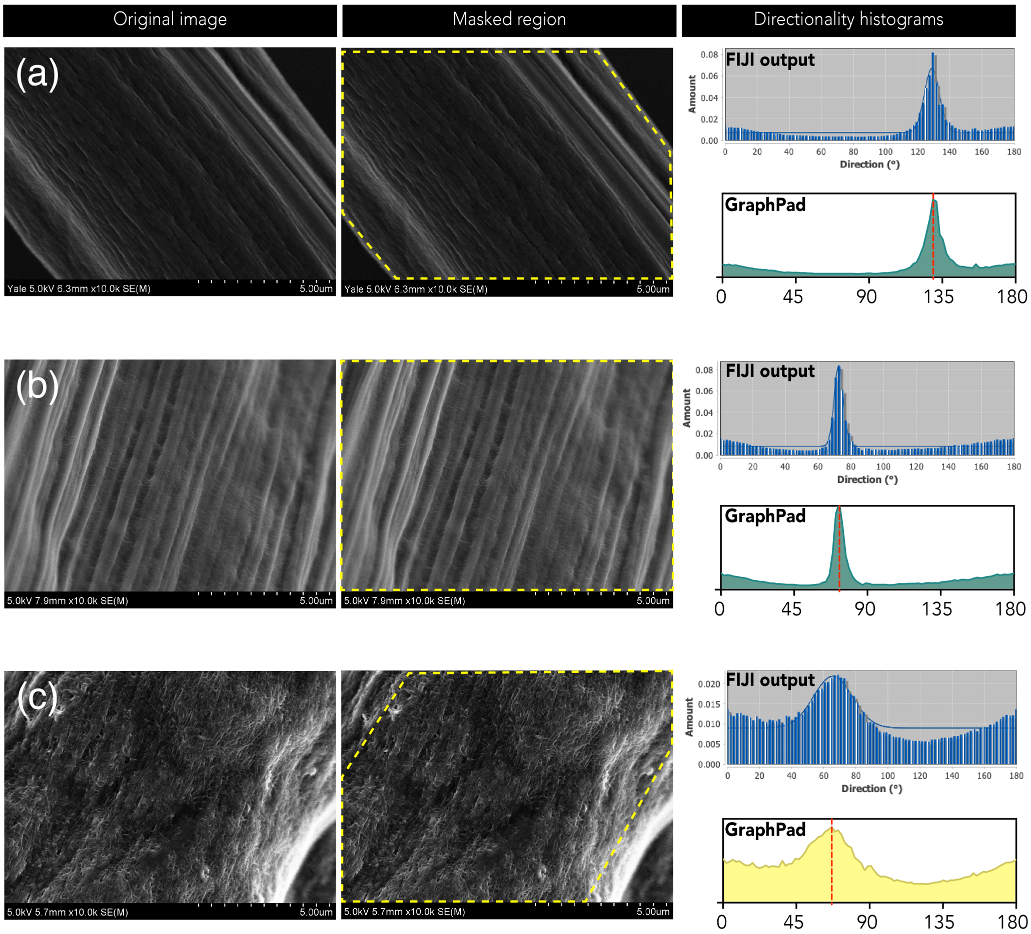


**Figure S3.** Image processing pipeline for quantitative analysis of feature orientation. This figure illustrates the step-by-step methodology used to quantify alignment from scanning electron micrographs. (Left column) The process begins with the raw SEM micrograph as acquired from the microscope. (Central column) To ensure data accuracy and eliminate artifacts, a mask is manually created for each image. The mask isolates the in-focus regions of the filament surface while excluding the background, out-of-focus edges, and any over-exposed areas. (Right column) The masked image is then analyzed using the Directionality plugin in Fiji (ImageJ2). The top right panel shows the histogram output from the plugin, plotting frequency against orientation angle. For improved visualization and statistical analysis, the raw bin counts are exported and replotted in GraphPad Prism (bottom right panel). The red line indicates the mean of the orientation distribution. The standard deviation of this distribution is used as the quantitative metric for alignment dispersion, as reported in the main text.

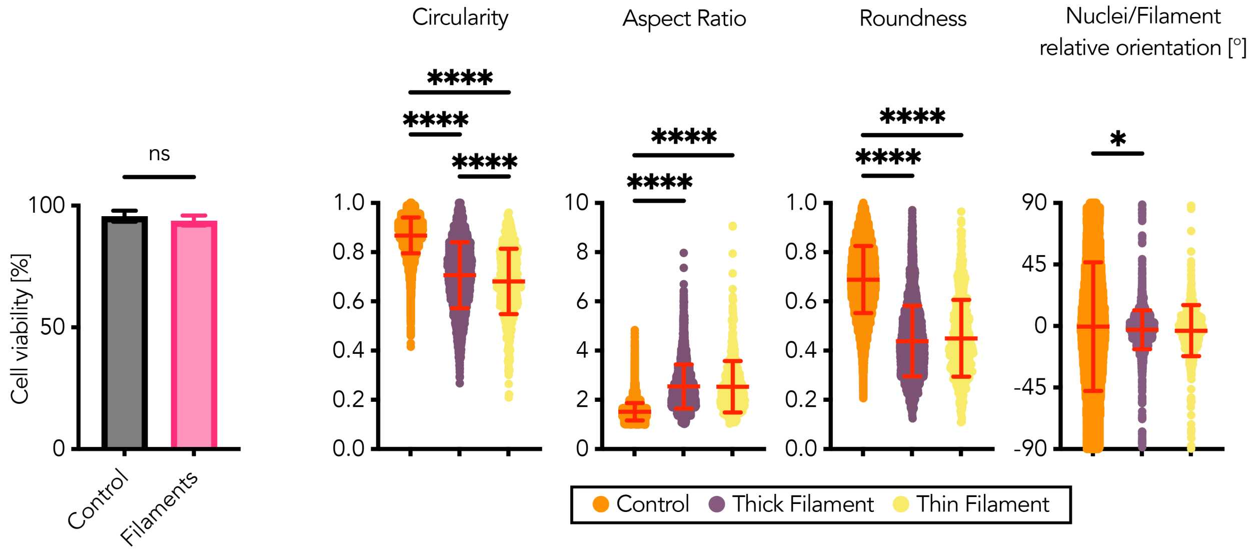


**Figure S4.** (Left Panel) Bar plot showing high cell viability (>95%) for mMSCs cultured on collagen I filaments compared to the glass control, with no statistically significant difference ("ns"), confirming the material's excellent biocompatibility. (Right Panels) Violin plots quantifying key morphological and orientation descriptors. The analysis demonstrates a profound topographical response: compared to cells on the 2D control, cell nuclei on filaments become significantly less circular and more elongated (higher aspect ratio). Just like cells, nuclei show strong alignment with the filament's major axis, in stark contrast to the disperse orientation observed on the 2D control. Results are presented as violin plots showing the distribution of all measurements, with the mean ± standard deviation overlaid. Asterisks (*) denote statistically significant differences (*$p < 0.05$, ****$p < 0.0001$).

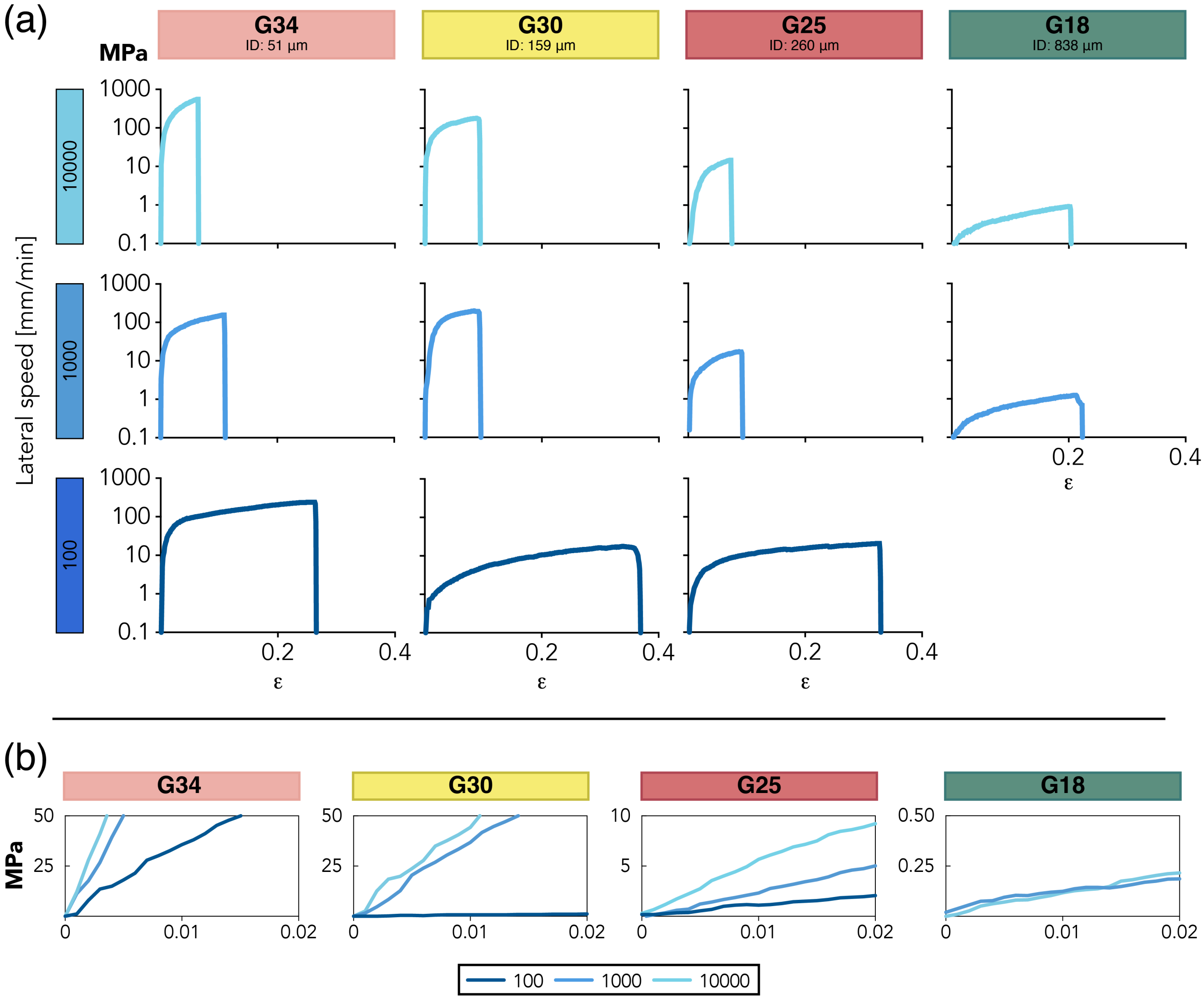


**Figure S5.** (a) Representative stress-strain curves for the tested fabrication parameters, grouped by nozzle size and color-coded by pulling speed. The stress axis is logarithmic to fit the wide range of material strengths, (b) Zoomed-in view of the region 0-2% strain used for the calculate Young's modulus for each of the tested conditions.

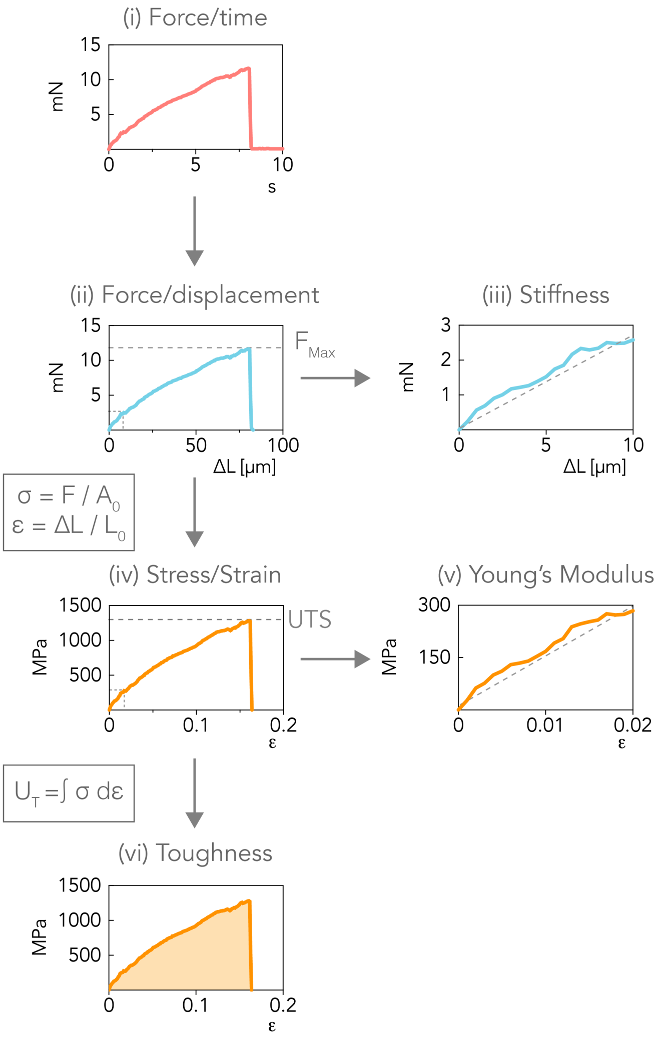


**Figure S6.** Data processing pipeline for the derivation of mechanical properties. This figure illustrates the complete workflow used to process raw tensile testing data and extract the reported mechanical metrics. (i) Raw force-time signal as acquired in LabVIEW. (ii) Using the known stretching rate of 0.1 mm/s, the time axis is converted to elongation ($\Delta L$) to generate a force-displacement curve. The rupture force ($F_{max}$) is identified from this plot. (iii) A linear regression is applied to the initial portion of the force-displacement curve (below 2% elongation); the slope of the liner fit equals the filament's Stiffness (N/m). (iv) Force and elongation data are normalised by the filament's initial cross-sectional area ($A_0$) and initial length ($L_0$), respectively, to obtain the engineering stress ($\sigma = F/A_0$) and engineering strain ($\epsilon = \Delta L/L_0$) curve. The peak of this curve represents the Ultimate Tensile Strength (UTS). (v) Similarly, a linear regression of the initial region of the stress-strain curve (below 2% strain) is used to determine the Young's Modulus (E). (vi) Finally, the Tensile Toughness ($U_T$) is calculated as the total energy absorbed per unit volume before fracture, it is visually represented by the area under the curve and determined by integrating the area under the entire stress-strain curve.

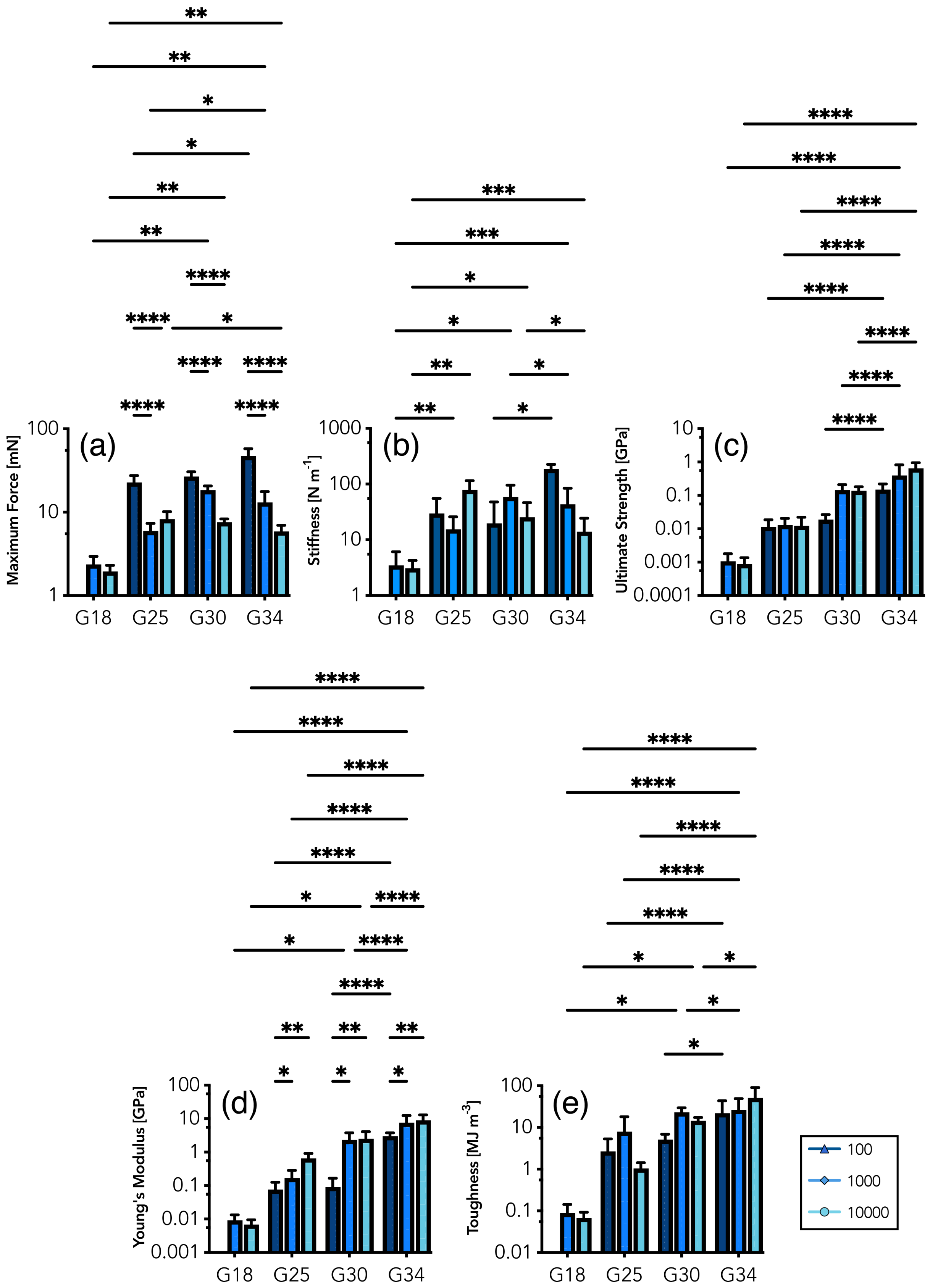


**Figure S7.** Mechanical properties of fabricated collagen filaments. Grouped bar plots showing the mean (± SD) for (a) Maximum Force (mN), (b) Stiffness (N/m), (c) Ultimate Tensile Strength (GPa), (d) Young's Modulus (GPa), and (e) Toughness (MJ/$m^3$). Results are grouped by nozzle size, with each group containing results for the three tested pulling speeds (100, 1000, 10000 mm/min). Each bar represents the mean value, and the error bars denote the standard deviation (SD) calculated from at least five independent filament tests ($n \geq 5$) per condition. Statistically significant differences between nozzle gauges were determined by a two-way ANOVA. Asterisks (*) denote statistical significance (* $p < 0.05$, ** $p < 0.01$, *** $p < 0.001$, ****$p < 0.0001$).
